# YTHDF1 and YTHDC1 m^6^A reader proteins regulate HTLV-1 *tax* and *hbz* activity

**DOI:** 10.1101/2024.11.18.624222

**Authors:** Emily M. King, Amanda Midkiff, Karsyn McClain, Sanggu Kim, Amanda R. Panfil

## Abstract

Human T-cell leukemia virus type 1 (HTLV-1) is a retrovirus responsible for adult T-cell leukemia/lymphoma (ATLL) and HTLV-1-associated myelopathy/tropical spastic paraparesis (HAM/TSP), a progressive neurodegenerative disease. Regulation of viral gene expression plays a key role in viral persistence and pathogenesis. However, the molecular mechanisms underlying this fine-tuned regulation remain poorly understood. Little is known regarding RNA chemical modifications of HTLV-1 RNA and how these affect viral biology and disease development. Posttranscriptional chemical modification of RNA is common in eukaryotes, with N^6^-methyladenosine (m^6^A) being the most prevalent. In this study, we investigated the role of m^6^A RNA modifications on HTLV-1 gene expression. Using MeRIP-Seq, we mapped sites of m^6^A modification to the 3’ end of the viral genome. We found HTLV-1 RNA, as well as viral oncogene transcripts *tax* and *hbz*, contained m^6^A modifications. m^6^A-depletion in HTLV-1-transformed cells decreased sense-derived viral genes (*Tax, Gag, Env*) and increased antisense-derived *Hbz* expression. *Tax* and *hbz* transcripts were bound by reader proteins YTHDF1 and YTHDC1 in a panel of HTLV-1 T-cell lines. Using expression vectors and shRNA-mediated knockdown, we found YTHDF1 had opposing effects on viral gene expression – decreasing sense-derived viral genes and increasing antisense-derived *Hbz*. Upon further study, the YTHDF1 effects on *tax* abundance were dependent on *tax* m^6^A deposition. The nuclear m^6^A reader protein YTHDC1 affected the abundance of both sense- and antisense-derived viral transcripts and specifically enhanced the nuclear export of *tax* transcript. Collectively, our results demonstrate global m^6^A levels and m^6^A reader proteins YTHDF1 and YTHDC1 regulate HTLV-1 gene expression.

**Importance:** Human T-cell leukemia virus type 1 (HTLV-1) persistence and pathogenesis are controlled through tight regulation of viral gene expression. The fate of RNA can be controlled by epigenetic modifications which impact gene expression without altering the DNA sequence. Our study details the impact of N6-methyladenosine (m^6^A) RNA chemical modifications on HTLV-1 gene expression. We found reductions in global m^6^A levels affected viral gene expression, decreasing *Tax* and other sense-derived viral genes, while increasing the antisense-derived *Hbz*. Our results suggest the oncogenic viral transcripts, *tax* and *hbz*, are m^6^A-modified in cells. We found these viral RNA modifications are interpreted by reader proteins YTHDF1 and YTHDC1, which dictate the fate of the viral RNA. Understanding HTLV-1 RNA chemical modifications offers potential insights into novel therapeutic strategies for HTLV-1-associated diseases.

## Introduction

Human T-cell leukemia virus type 1 (HTLV-1) is an oncogenic retrovirus endemic in Southwestern Japan, sub-Saharan Africa, South American and the Caribbean, with foci throughout the Middle East and Australia (1, 2). Approximately 5-10 million people worldwide are infected with HTLV-1 (1). However, this number is likely an underestimate given the modes of transmission (mother-to-child through breastfeeding, sexual contact, exposure to infected blood products), lack of large epidemiologic studies, and insufficient screening/prevention in many countries (2–4). HTLV-1 is the causative agent of two main diseases: adult T-cell leukemia/lymphoma (ATLL), a chemotherapy-resistant non-Hodgkin’s malignancy of CD4+ T-cells, and HTLV-1-associated myelopathy/tropical spastic paraparesis (HAM/TSP), a chronic, progressive neurogenerative disease (5). HTLV-1 infection has also been implicated in several inflammatory diseases, including uveitis, keratitis, conjunctivitis and dermatitis (6). Diseases caused by HTLV-1 often have a clinical latency period of several decades after infection.

HTLV-1 is a complex retrovirus that encodes regulatory and accessory genes, which induce transformation and stimulate cellular proliferation, resulting in viral persistence (7–9). The sense strand of the integrated proviral genome encodes structural and enzymatic genes, *Gag, Pro, Pol* and *Env*, that are typical of all retroviruses. The sense strand also encodes regulatory genes *Tax* and *Rex,* and accessory genes *p12/p8, p13,* and *p30* (10). HTLV-1 also encodes an accessory/regulatory gene within the antisense strand of the genome, termed *Hbz* (11, 12). Early proviral transcription and cellular transformation are driven by Tax through its interaction with several important signaling pathways involved in cell survival, proliferation, and genomic stability, such as persistent activation of NF-κB (13). Conversely, one of the many roles of Hbz is to regulate Tax-mediated activation of viral transcription (14–17). Through interactions with cellular transcription factors (ATF-1, CREB, JunB, JunD, c-Jun) via its basic leucine zipper domain, Hbz can sequester these factors away from DNA and inhibit Tax-mediated transcription (18). Hbz can also antagonize other roles of Tax, including stimulation of the classical NF-κB pathway (19). By regulating the activity of Tax, Hbz promotes survival of HTLV-1-infected CD4+ T-cells via downregulation of NF-κB, controlled proliferation, and clonal expansion. Moreover, Hbz-deleted virus has decreased persistence and disease development *in vivo* (20), further highlighting its importance in infection and oncogenesis.

N-6-methyladenosine (m^6^A) represents methylation of the N6 position of adenine. As the most abundant post-transcriptional chemical modification in eukaryotic RNA, it has several documented roles in viral gene regulation and pathogenesis (21). m^6^A modifications are enriched near stop codons and 3’ untranslated regions (UTRs) (22–24), suggestive of a critical role in cells. Typically m^6^A is preferentially deposited on “DRACH” motifs (D = G/A/U, R = G>A, and H = U>A>C) (25). This dynamic modification is deposited onto RNA transcripts by m^6^A “writers,” removed by “erasers,” and interpreted by “readers.” The writer proteins are responsible for transferring a methyl group from an S-Adenysyl Methionine (SAM) molecule onto the 6^th^ nitrogen of an adenosine base (26). METTL3 and METTL14 are two currently identified writers in mammalian cells. Together, these writers form the m^6^A-METTL complex (MAC) and introduce m^6^A into nascent RNA transcripts (22, 23, 27, 28). MAC interacts with the m^6^A-METTL-associated complex (MACOM), which is composed of proteins with various functions that aid in the stability, binding, and methylation, including Wilm’s Tumor Associated Protein (WTAP), Zinc Finger CCCH-Type Containing 13 (ZC3H13), RNA-Binding Motif Protein 15B (RBM15/15B), Vir Like m^6^A Methyltransferase Associated (VIRMA) and E3 Ubiquitin-Protein Ligase Hakai (HAKAI).

The m^6^A eraser proteins are AlkB homolog 5 (ALKBH5) and fat mass and obesity associated protein (FTO). The function of these proteins is facilitated through enzymes in the alpha-ketoglutarate-dependent dioxygenase (AlkB) family, in which they utilize the alpha-KG domain to recognize the m^6^A modified RNA and bind Fe(II) and alpha-KG cofactors. Upon substrate recognition, the demethylases catalyze oxidative phosphorylation of m^6^A to regenerate an unmodified adenosine (29–32). RNA transcripts modified by m^6^A are interpreted by “readers” (YTHDF1-3, YTHDC1-2) (19, 33–37), members of the YTH domain-containing family of proteins. YTHDF1 has been shown to regulate 5’ cap-dependent translation by interacting with the 5’ UTR-associated eIF3 protein (34). Additionally, m^6^A modification in 5’ UTRs can facilitate 5’ cap-independent translation by directly recruiting eIF3 (38). YTHDF2 regulates mRNA stability by recruiting the Carbon Catabolite Repressive – Negative on TATA-less (CCR4-NOT) deadenylase complex to promote degradation of m^6^A-modified RNA(33, 37). YTHDF3 has overlapping functions with both YTHDF1 and YTHDF2 and can alter both translation and decay, however the exact mechanistic role is yet to be elucidated (36, 39). Functional roles of these cytoplasmic readers, however, remain partially controversial (40–42), potentially reflecting their context-dependent roles. YTHDC1 is the only reader protein localized to the nucleus, promoting splicing and nuclear export of m^6^A-modified pre-mRNAs through interaction with splicing factors SRSF3 and SRSF10, the former of which interacts with nuclear RNA export factor 1 (NXF1) to facilitate export (35). The function of YTHDC2 has not been fully elucidated in the context of m^6^A.

In this study, we identify, map, and characterize m^6^A RNA modification in HTLV-1, demonstrating the versatile role this post-transcriptional modification plays in viral transcript abundance and localization. A better understanding of the chemical modifications on HTLV-1 RNA and how m^6^A regulatory machineries control *tax* and *hbz* expression is warranted. These studies will help uncover potential therapeutic targets, paving the way for novel treatment strategies against HTLV-1 and associated diseases.

## Materials and Methods

### Cell culture

All cell lines were cultured in media containing 10% fetal bovine serum (FBS), 100 U/mL penicillin, 100 μg/mL streptomycin, and 2 mM L-glutamine and maintained in a humidified atmosphere of 5% CO2 and air at 37°C, unless otherwise noted. Human embryonic kidney (HEK) 293T cells were cultured in Dulbecco’s modified Eagle’s medium (DMEM) (Thermo Fisher Scientific, Waltham, MA). SLB-1 (HTLV-1-transformed T-cell line) cells were cultured in Iscove’s DMEM (Mediatech, Inc. Manassas, VA). ATL-ED (ATLL patient-derived human T-cell line), HuT-102 (HTLV-1-transformed T-cell line), C91/PL (HTLV-1-transformed T-cell line) and JET cells (Jurkat indicator cells expressing tdTomato under the control of 5 times tandem repeat of Tax responsive element [TRE]) (43–45) were cultured in Roswell Park Memorial Institute (RPMI) 1640 medium (Thermo Fisher Scientific). HTLV-1-immortalized primary human T-cell lines (PBL-HTLV-1) were cultured in RPMI 1640 supplemented with 20% FBS and 10 U/mL recombinant human interleukin-2 (hIL-2; Roche Diagnostics GmbH, Mannheim, Germany).

### Plasmids

The pPB-CAG plasmid vector was used to express YTHDF1, as previously described (46). The pPB-CAG plasmid vector was used to express YTHDC1 (kind gift from Dr. Li Wu, University of Iowa). The plasmid vector used to express native *tax* sequence (pCMV-Tax) was generated as previously described (47). The plasmid vector used to express the altered *tax* sequence was designed using the HTLV-1 ACH molecular clone as reference sequence (48). After human codon optimization, the Tax sequence was cloned into vector pcDNA3.1(+) at the BamHI/NheI restriction enzyme sites (GenScript; Piscataway, New Jersey) to produce the altered Tax sequence expression vector (pCMV- ALT. Tax). The plasmid containing the wildtype HTLV-1 infectious proviral clone, ACHneo, was described previously (49). The LTR-1-lucuferase and TK- renilla reporter plasmids were described previously (50).

### FTO demethylation of HTLV-1 RNA

Demethylation of HTLV-1 RNA m^6^A sites was performed as previously described (51). Briefly, viral particles produced by PBL-HTLV-1 cells were concentrated using ultracentrifugation in a Sorvall SW-41 swinging bucket rotor and resuspended in TRIzol (Thermo Fischer Scientific). RNA was isolated following the manufacturer’s protocol (Invitrogen, Carlsbad, CA). RNA was subsequently enriched for mRNA using polyA bead capture (Qiagen, Redwood City, CA) to exclude other m^6^A-modified RNAs, such as tRNA. 500ng of mRNA was used for FTO treatment in a 100 µL reaction buffer containing 50 mM HEPES buffer (pH 7.0), 75 µM (NH4)_2_Fe (SO4)_2_•6H_2_O, 2 mM L-ascorbic acid, 300 µM alpha-ketoglutaric acid, 200 U RNAsin ribonuclease inhibitor, 5 µg/mL BSA, and titrating amounts (0.1 µM, 0.3 µM, 0.6 µM) of recombinant FTO protein (Abcam, Cambridge, MA). The reaction mixture was incubated at 37°C for 1 hour and the reaction was stopped with 5 mM EDTA. RNA samples (treated with or without FTO) were immediately applied to an m^6^A ELISA (Abcam) for m^6^A quantification according to the manufacturer’s protocol.

### m^6^A depletion using STM2457

STM2457 (Sigma-Aldrich, St. Louis, MO; CAS#2499663-01-1), a selective METTL3 inhibitor, was used to prevent m^6^A deposition. 5 x 10^5^ C91/PL, HuT-102, SLB-1, ATL-ED, PBL-HTLV-1, or HEK293T cells were seeded in a 6-well dish and incubated at 37°C with 60 µM of STM2457 for 72 hours. Cell viability was measured using Trypan Blue staining. Cells were collected for downstream experiments (p19 Gag ELISA, JET infectivity assay) and analysis (western blot, RT-qPCR). Successful depletion of m^6^A in total RNA was confirmed using an m^6^A ELISA (Abcam), following the manufacturer’s protocol.

### MeRIP-Seq

Methylated RNA immunoprecipitation sequencing (MeRIP-Seq) was performed as previously described (52). Briefly, total RNA from PBL-HTLV-1 or SLB-1 cells was extracted using TRIzol (Thermo Fischer Scientific). Poly (A) RNA was purified from 50 μg total RNA using Dynabeads Oligo (dT)25-61005 (Thermo Fisher Scientific) using two rounds of purification. Enriched poly(A) RNA was fragmented into small pieces using Magnesium RNA Fragmentation Module (New England Biolabs, cat.e6150, USA) under 86℃ for 7 m. The cleaved RNA fragments were incubated for 2 h at 4℃ with m^6^A-specific antibody (No. 202003, Synaptic Systems, Germany) in IP buffer (50 mM Tris-HCl, 750 mM NaCl and 0.5% Igepal CA-630). IP RNA was reverse-transcribed to create the cDNA by SuperScript™ II Reverse Transcriptase (Invitrogen, cat. 1896649, USA), followed by synthesis of U-labeled second-stranded DNAs with E. coli DNA polymerase I (NEB, cat.m0209), RNase H (NEB, cat.m0297), and dUTP Solution (Thermo Fisher Scientific, Cat. R0133). A-tailing was added to the blunt ends of each strand, preparing them for ligation to indexed adapters. Single- or dual-index adapters were ligated to the fragments, and size selection was performed with AMPureXP beads. After heat-labile UDG enzyme (NEB, Cat. m0280) treatment of the U-labeled second-stranded DNAs, the ligated products were amplified with PCR by the following conditions: initial denaturation at 95℃ for 3 m; 8 cycles of denaturation at 98℃ for 15 sec, annealing at 60℃ for 15 sec, and extension at 72℃ for 30 sec; and final extension at 72℃ for 5 m. The average insert size for the final cDNA library was 300 ± 50 bp. Sequencing was performed with 2 × 150bp paired-end (PE150) mode on Illumina Novaseq™ 6000 following the vendor’s recommended protocol. Data was analyzed using Integrated Genome Browser (http://www.igv.org). Input and raw sequencing data was visualized against an HTLV-1 reference genome (ACHneo) to visualize peaks of m^6^A enrichment. Generated IGV peaks were created by normalizing the IP sample peaks to the input sample peaks. Mapped reads of IP and input libraries were input for R package *exomePeak* (https://bioconductor.org/packages/exomePeak) for identification of significant m^6^A peaks with bed or bigwig format that can be adapted for visualization with IGV software. Differential expression of peak was selected with a *p* value < 0.05 by R package edgeR (https://bioconductor.org/packages/edgeR).

### RNA crosslinking and immunoprecipitation

10^7^ vehicle (DMSO) or STM2457-treated C91/PL, PBL-HTLV-1, SLB-1, and ATL-ED cells were collected and washed with ice-cold 1X PBS. Cells used for reader protein cross-linking were UV-crosslinked using 1500J, three times. The cells were lysed in 1mL of NP-40 lysis buffer containing protease inhibitor cocktail and 2000 U of RNAse inhibitor. The solution was rocked for 30 minutes and centrifuged at 13,000 x *g* for 10 minutes at 4°C. 50 µL of sample was saved as input, while the remaining supernatant was evenly divided for immunoprecipitation using 3 µg of antibody: IgG-rabbit, m^6^A (Abcam, ab151230), YTHDF1 (Abcam, ab220162), YTHDF2 (Abcam, ab220163), YTHDF3 (Abcam, ab220161), or YTHDC1 (Abcam, ab264375). Supernatant and antibody were incubated together overnight at 4°C with rocking. Protein G Dyanabeads™ (Thermo Fischer Scientific) were added to the antibody-complexed samples and rocked at 4°C for 2 hours. Bead complexes were then washed three times using 1mL of NP-40 lysis buffer. 250 µL of TRIzol (Thermo Fischer Scientific) was added to both the input and immunoprecipitated samples for RNA isolation following the manufacturer’s protocol. A portion of sample containing non-immunoprecipitated and immunoprecipitated protein lysate was saved for western blot to confirm successful reader protein immunoprecipitation.

### Lentiviral production and cell transduction

Lentiviral vector expressing short hairpin RNA (shRNA) targeting YTHDF1 or YTHDC1 was previously described (46, 53). The universal negative control, pLKO.1 (RHS4080) was purchased from Open Biosystems (Dharmacon, LaFayette, CO, USA) and propagated according to the manufacturer’s instructions. HEK293T cells were transfected with lentiviral vector(s), as well as DNA vectors encoding HIV Gag/Pol and vesicular stomatitis virus G in 10 cm dishes using Lipofectamine® 2000 reagent according to the manufacturer’s instructions. 72 hours following transfection, media containing lentiviral particles was collected and concentrated using ultracentrifugation in a Sorvall SW-41 swinging bucket rotor at 21,000 x *g* for 1.5h at 4°C. DMSO- or STM2457-treated target cells (SLB-1, ATL-ED) were transduced using polybrene (8 µg/mL) and spin-inoculation at 2000 x *g* for 2 hours at room temperature. 72 hours post-transduction, cells were selected using 1-2 µg/mL puromycin for 3 days and subsequently harvested for downstream experiments (p19 Gag ELISA) or analysis (RT-qPCR, western blot).

### Plasmid transfection

DMSO- or STM2457-treated HEK293T cells were co-transfected with 100 ng YTHDF1, 1 µg ACHneo, 100 ng LTR-1-luc, and 20 ng rtk-luc, where indicated, using Lipofectamine™ 2000 Transfection Reagent (Invitrogen). 72 hours post-transfection, cells were harvested for downstream analysis. For analysis of YTHDF1 and YTHDC1 on Tax protein, DMSO- or STM2457-treated HEK293T cells were co-transfected with titrating amounts (0 ng, 10 ng, 50 ng, or 100 ng) of YTHDC1 or YTHDF1 and 500 ng of pCMV-Tax (native *tax* sequence) or pCMV- ALT. Tax (altered *tax* sequence) using Lipofectamine™ 2000 Transfection Reagent (Invitrogen) following the manufacturer’s protocol. Cells were collected 48 hours post-transfection for downstream RNA fractionation and western blot analysis.

### RNA fractionation

HEK293T cells were collected and subjected to nuclear/cytoplasmic fractionation using a previously established protocol (54). Cytoplasmic and nuclear fractions were suspended in TRIzol (Invitrogen) for RNA extraction following the manufacturer’s protocol and used for subsequent RT-qPCR analysis. 2 µL of RNA was used for cDNA synthesis using the SuperScript IV First-Strand Synthesis System (Invitrogen). qPCR was performed as described below in ‘Quantitative RT-PCR’. Lysate derived from the fractionated samples was applied to a western blot and immunoblotted with the cytoplasmic protein β-tubulin or the nuclear protein Lamin A/C to confirm successful fractionation.

### Luciferase reporter assays

DMSO- or STM2457-treated HEK293T cells were transfected with YTHDF1, ACHneo, LTR-1- luciferase and TK-renilla. Cells were collected after 48 h and lysed in Passive Lysis Buffer (Promega, Madison, WI). Relative firefly and Renilla luciferase units were measured using a FilterMax F5 MultiMode Microplate Reader (Molecular Devices, San Jose, CA) using the Dual-Lufierase® Reporter Assay System (Promega, Madison, WI) according to the manufacturer’s instructions. Each condition was performed in triplicate.

### p19 Gag ELISA

Cells were seeded at 2.5 x 10^5^/mL in a 6-well plate at 37°C. 24 hours later, supernatant was collected to measure HTLV-1 p19 Gag production using the RETROTEK HTLV p19 Antigen ELISA (ZeptoMetrix Corporation, Buffalo, NY) according to the manufacturer’s instructions.

### JET Infectivity Assay

1.0 x 10^4^ STM2457- or DMSO-treated C91/PL cells were co-cultured with 5.0 x 10^4^ JET cells in a 12-well plate at 37°C. 48 hours later, cells were collected by centrifugation and TdTomato expression was quantified using flow cytometry.

### Quantitative RT-PCR

Total RNA was isolated from 1 x 10^6^ cells per condition using TRIzol (Invitrogen) according to the manufacturer’s instructions. Isolated RNA was quantitated using the ND-1000 Nanodrop spectrophotometer (Thermo Fisher Scientific) and DNAse-treated using recombinant DNAse I (Sigma-Aldrich, Cat#04716728001). 200 ng of total RNA was used for cDNA synthesis using the SuperScript IV First-Strand Synthesis System (Invitrogen). 2 µL of cDNA was used per qPCR reaction with iQ SYBR Green Supermix (Bio-Rad, Hercules, CA) and 300 nM of each sense and antisense primer (20 µL total reaction volume). Reactions were performed in 96-well plates using the CFX96 Touch Real-Time PCR Detection System (Bio-Rad). The reaction conditions were 50°C for 2 min, 95°C for 10 min, followed by 40 cycles of 15 sec at 95°C and 1 min at 60°C. Primer pairs to detect viral mRNA species (*tax, gag, env, hbz*) and *gapdh* were described previously(55, 56).

### Immunoblotting

Total protein in cell lysates was quantitated using the Pierce™ BCA Protein Assay Kit (Thermo Fisher Scientific) and FilterMax F5 Multi-Mode Microplate Reader (Molecular Devices). Protein was loaded in equal amounts on 4 – 20% Mini-PROTEAN® TGX™ Precast Protein Gels (Bio-Rad) and transferred onto Amersham™Protran® Western blotting nitrocellulose membranes (Cytiva, Marlborough, MA). Membranes were blocked with 5% milk in 1x PBS with 0.1% Tween-20 and incubated with the following primary antibodies: YTHDF1 (1:1000; Abcam, ab220162), YTHDF2 (1:1000; Abcam, ab220163), YTHDF3 (1:1000; Abcam, ab220161), YTHDC1 (1:10,000; Abcam, ab264375), Tax (rabbit anti-sera), HBZ (rabbit anti-sera) (57), gp46 (1:1000; Santa Cruz Biotechnology, sc-53890), p19/gag (1:1000; ZeptoMetrix SKU0801116), SP1 (1:1000; Cell Signaling #5931), JunD (1:1000; Santra Cruz Biotechnology, sc-271938), MEF-2A (1:1000, Proteintech, 12382-1-AP), MEF-2C (1:1000, Proteintech, 10056-1-AP), β-tubulin (1:1000; Cell Signaling #2146), Lamin A/C (1:1000; Cell Signaling #4777), and β-actin (1:5000; Sigma-Aldrich, A2228). Membranes were developed using Pierce™ ECL Western Blotting Substrate (Thermo Fisher Scientific) and captured using an Amersham Imager 600 (GE Healthcare Life Sciences).

### Statistics

Statistical analyses were performed using GraphPad Prism 7 software (GraphPad Software) as indicated. Studies were analyzed by the Student’s t-test. Statistical significance was defined as * p ≤ 0.05, ** p ≤ 0.01, *** p ≤ 0.001, and **** p ≤ 0.0001. Image J was used to quantify immunoblots. Each experiment was performed in triplicate, with representative immunoblots shown in each figure.

## Results

### HTLV-1 RNA is m^6^A-modified

RNA viruses, including HTLV-1, maximize the utility of every feature within their compact RNA genomes (58–62). To date, over 300 types of RNA chemical modifications (collectively termed the ‘epitranscriptome’) have been identified (63, 64), introducing a new layer of regulation in a range of viral processes and cancers (64–68). To determine whether m^6^A modifications are present on the HTLV-1 RNA genome, virion RNA from purified viral particles was quantified using an m^6^A ELISA (**Figure 1A**). Treatment of the RNA with recombinant FTO (m^6^A eraser) resulted in a dose-dependent decrease of m^6^A-modified RNA. Mapping of m^6^A-modified regions within the HTLV-1 RNA genome was performed using methylated RNA immunoprecipitation and sequencing (MeRIP-Seq) in an HTLV-1 newly immortalized T-cell line (PBL-HTLV-1) and an HTLV-1-transformed T-cell line (SLB-1). A consistent m^6^A enrichment peak was generated near the regulatory pX region (3’ end of the genome) which encodes both *tax* and *hbz* (**Figure 1B**). Notably, a second peak was inconsistently generated near the 2kb region of the RNA, which may correspond to m^6^A enrichment in the Gag or Pro genes. These results were supported through m^6^A RNA immunoprecipitation in the HTLV-1-transformed T-cell lines SLB-1 and C91/PL, as well as the ATLL patient-derived T-cell line, ATL-ED. We found m^6^A modification of *tax* (**Figure 1C**), *gag* (**Figure 1D**), *env* (**Figure 1E**) and *hbz* (**Figure 1F**) in SLB-1, C91/PL and ATL-ED (expresses *hbz* only) cell lines. m^6^A enrichment was abolished upon treatment of cells with a selective METTL3 inhibitor (STM2457). Successful decrease of total m^6^A-modified RNA was confirmed in each cell line using an m^6^A ELISA (**Supplemental Figure 1A**). Importantly, STM2457 did not significantly affect cell viability during the duration of our experiments (**Supplemental Figure 1B**). Taken together, these data indicate the presence of m^6^A modifications on viral genomic RNA, as well as viral sense (*tax, gag, env*) and antisense (*hbz*) transcripts in T cells

**Figure 1.**
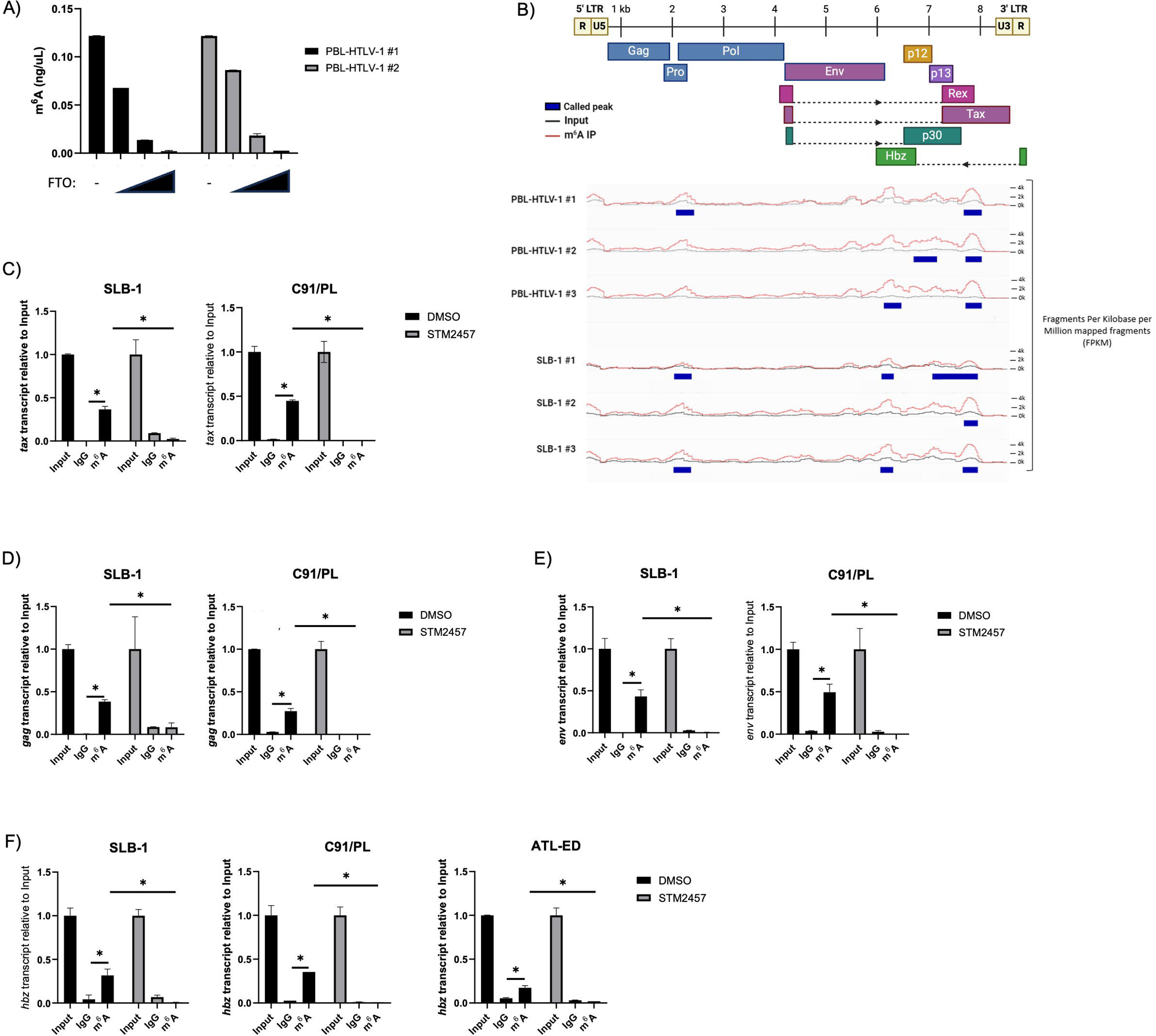
HTLV-1 RNA contains m^6^A modifications. A) mRNA was isolated from virus produced by two different newly immortalized PBL-HTLV-1 T-cell lines. Portions of the mRNA samples were treated with titrating amounts (0.1 µM, 0.3 µM, 0.6 µM) of recombinant FTO eraser protein. The level of m^6^A-modified mRNA was quantified in vehicle and FTO-treated conditions using an ELISA. B) MeRIP-Seq was performed on RNA isolated from PBL-HTLV-1 and SLB-1 T-cell lines. Sites of m^6^A enrichment were aligned with the viral RNA genome to generate peaks of m^6^A deposition using Integrated Genome Viewer. PBL-HTLV #1 peaks: 2130-2468, 7632-7975; PBL-HTLV #2 peaks: 6668-7120, 7663-7972; PBL-HTLV #3 peaks: 6106-6433, 7669-7975; SLB-1 #1 peaks: 2149-2482, 6107-6343, 7074-7938; SLB-1 #2 peak: 7675-7938; SLB-1 #3 peaks: 2134-2471, 6079-6322, 7657-7940. SLB-1, C91/PL and ATL-ED cell lines were treated with 60 μM STM2457 for 72 h and subjected to RNA immunoprecipitation using a m^6^A or IgG control antibody. The level of C) *tax* D) *gag* E) *env* and F) *hbz* transcripts was measured using qRT-PCR. Statistical significance was determined using a Student’s *t*-test; \**p* ≤ 0.05.

### Cellular m^6^A levels regulate viral gene expression

The effect of global m^6^A-level changes was assessed in C91/PL cells using titrating amounts of STM2457. A dose-dependent decrease in m^6^A-modified RNA was confirmed by m^6^A ELISA (**Figure 2A**). A decrease in cellular m^6^A resulted in dose-dependent decreases in sense (Tax, Gag, Env) protein (**Figure 2B**) and transcript levels (**Figure 2E**). Interestingly, however, it resulted in dose-dependent increases in antisense (Hbz) proteins and transcripts. As expected, based on the level of sense-derived gene products, the amount of p19 (Gag) released in the supernatant (**Figure 2C**) and the amount of infectious viral particles produced in C91/PL cells (**Figure 2D**) was decreased in response to STM2457 treatment. Importantly, we found similar decreases in sense (Tax, Gag, Env) and increases in antisense (Hbz) protein (**Figure 2G**) and transcript levels (**Figure 2F**) in HTLV-1-transformed SLB-1 and Hut-102 T-cell lines treated with STM2457. Similarly, STM2457 treatment decreased the amount of p19 (Gag) released into the supernatant in SLB-1 and Hut-102 cells (**Figure 2H**). Together, these results suggest global m^6^A levels regulate the differential gene expression between sense and antisense-derived viral genes.

**Figure 2.**
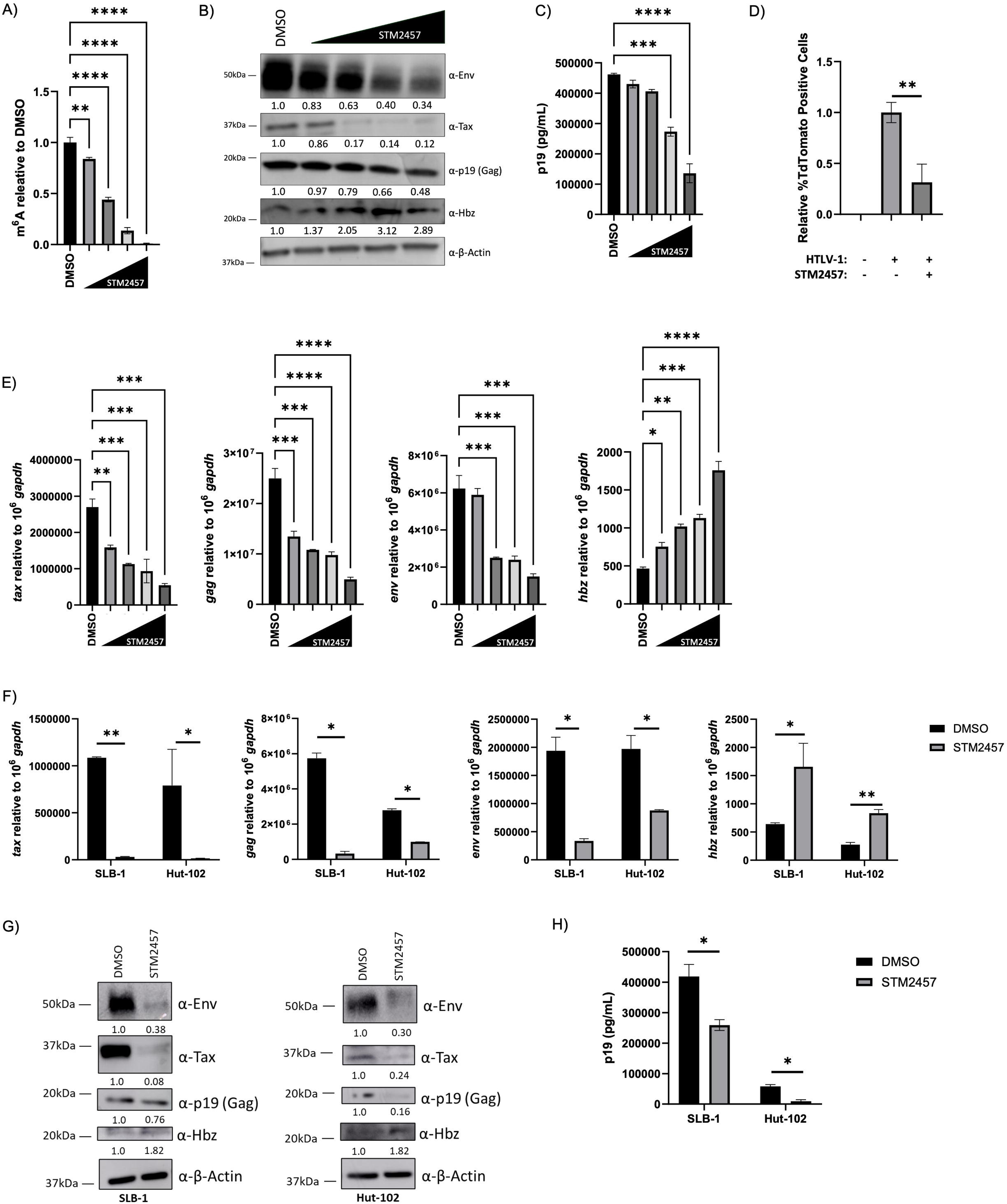
Cellular m^6^A depletion decreases sense-derived viral genes and increases *hbz* expression. C91/PL (HTLV-1-transformed T-cell line) cells were treated with vehicle or titrating amounts (20 μM, 40 μM, 60 μM, 80 μM) of STM2457 for 72 h. A) The level of m^6^A-modified mRNA in the cell was quantified using an ELISA. B) Western blot analysis was used to measure viral protein (Env, Tax, p19 [Gag], Hbz) expression. β-actin was used as a loading control. Relative band intensity was quantified using ImageJ. Protein quantification relative to DMSO (normalized to β-actin) is depicted below each western blot. C) Virus release in the cell supernatant was measured using a p19 Gag ELISA. D) Infectious virus release was quantified by measuring TdTomato expression in JET cells through flow cytometry. E) RNA was isolated and qRT-PCR was used to measure sense-derived transcripts (*tax, gag,* and *env*) and antisense-derived (*hbz*) transcripts. HTLV-1-transformed T-cell lines (SLB-1, Hut-102) were treated with or without 60 μM STM2457 for 72 h. F) RNA was isolated and qRT-PCR was used to measure sense-derived transcripts (*tax, gag,* and *env*) and antisense-derived (*hbz*) transcripts. G) Western blot analysis was used to measure viral protein (Env, Tax, p19 [Gag], Hbz) expression. β-actin was used as a loading control. Relative band intensity was quantified using ImageJ. Protein quantification relative to DMSO (normalized to β-actin) is depicted below each western blot. H) Virus release in the cell supernatant was measured using a p19 Gag ELISA. Statistical significance was determined using a Student’s *t*-test; \**p* ≤ 0.05, \*\**p* ≤ 0.01, \*\*\**p* ≤ 0.001, ****p≤ 0.0001.

### Reader proteins YTHDF1 and YTHDC1 bind *tax* and *hbz* RNA

Since both *tax* and *hbz* transcripts were identified as m^6^A-modified, we next sought to identify which m^6^A reader proteins (YTHDF1-3, YTHDC1) bind these transcripts. RNA cross-linked immunoprecipitation (RNA CLIP) was performed in vehicle (DMSO) or STM2457-treated SLB-1, PBL-HTLV-1, and ATL-ED cell lines. ATL-ED are an ATLL patient-derived T-cell line that does not produce sense transcripts (only antisense-derived *hbz*) due to hypermethylation at the 5’ LTR. Successful immunoprecipitation of each reader protein was confirmed by western blot (**Supplemental Figure 2A**). Using qRT-PCR, we found YTHDF1 and YTHDC1 consistently bound *tax* (**Figure 3A**) in PBL-HTLV-1 and SLB-1 T-cell lines and *hbz* (**Figure 3B**) in all cell lines. Furthermore, this binding is dependent on m^6^A modification as the binding was abolished in STM2457-treated cells. Interestingly, there was significant binding of YTHDF3 to *tax* and *hbz* in several cell lines and YTHDF2 to *hbz* in ATL-ED cells. However, we chose to focus on YTHDF1 and YTHDC1 which consistently bound *tax* and/or *hbz* across several different HTLV-1 cell lines. To ensure the m^6^A reader proteins were adequately expressed in the tested cell lines, we also confirmed expression of each reader protein (YTHDF1-3, YTHDC1) by western blot (**Supplemental Figure 2B**).

**Figure 3.**
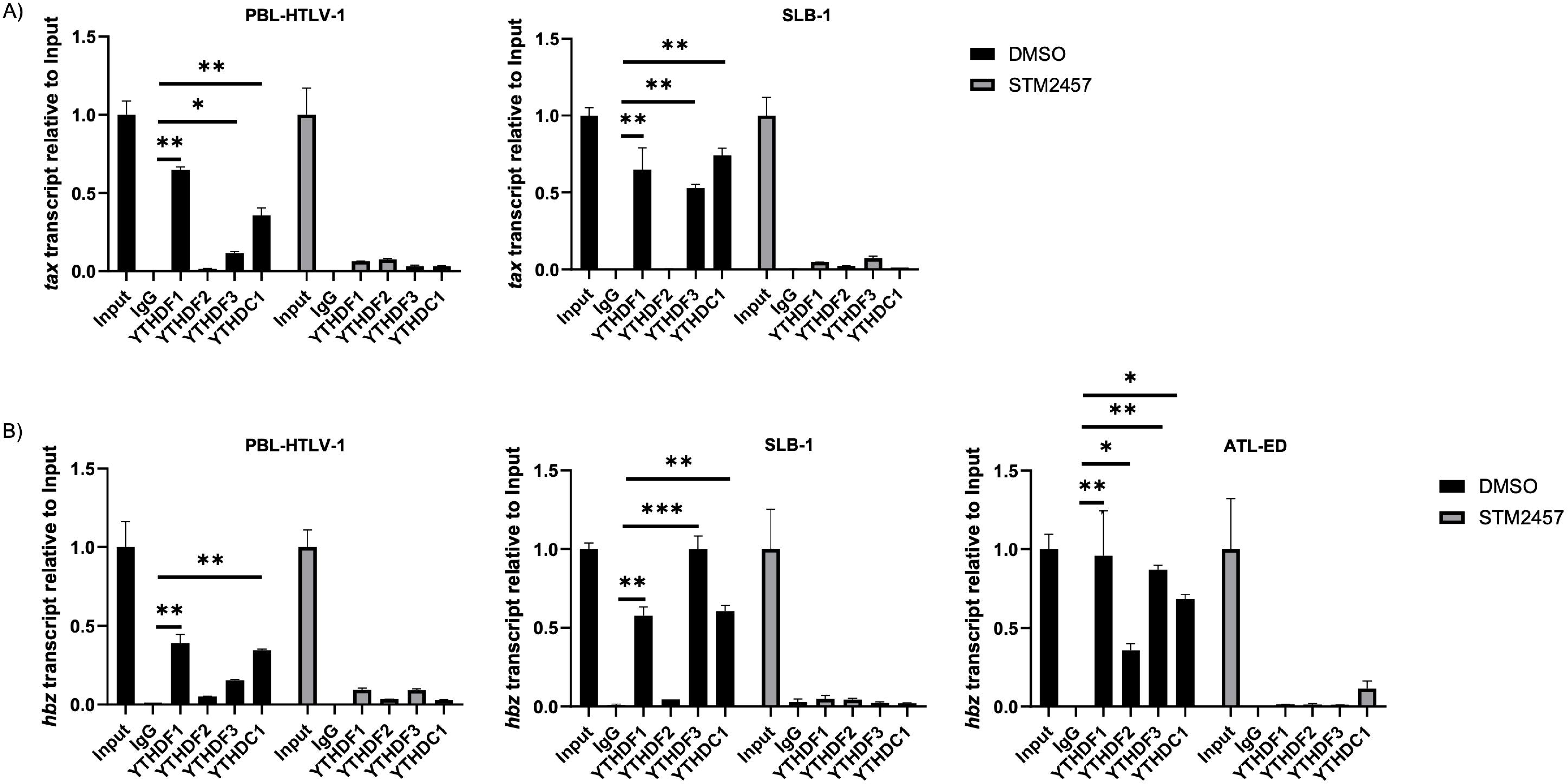
m^6^A reader proteins YTHDF1 and YTHDC1 bind *tax* and *hbz*. HTLV-1-transformed (SLB-1), HTLV-1-newly immortalized (PBL-HTLV-1) and ATL patient-derived (ATL-ED) T-cell lines were treated with or without 60 μM STM2457 for 72 h and subjected to RNA cross-linking and immunoprecipitation using antibodies against YTHDF1-3, YTHDC1 and an IgG control. Total RNA was isolated by TRIzol extraction and qRT-PCR was used to measure A) *tax* and B) *hbz* transcript abundance relative to Input. Statistical significance was determined using a Student’s *t*-test; \**p* ≤ 0.05, \*\**p* ≤ 0.01, \*\*\**p* ≤ 0.001.

### YTHDF1 differentially regulates sense and antisense-derived viral transcripts

To further dissect the role of m^6^A reader protein YTHDF1 in the context of HTLV-1, YTHDF1 was expressed in vehicle (DMSO) or STM2457-treated HEK293T cells in the presence of HTLV-1 provirus. Using an LTR-driven luciferase reporter vector (activated by Tax expression), we found expression of YTHDF1 significantly decreased viral sense transcription approximately 5-fold (**Figure 4A**). Pharmacological inhibition of m^6^A deposition (STM2457 treatment) abolished the YTHDF1-mediated transcriptional repression. Using qRT-PCR, we found YTHDF1 expression decreased sense-derived viral transcripts (*tax, gag, env*), while it increased antisense-derived viral transcript *(hbz*) **(Figure 4B**). Consistent with viral transcript alterations, the level of viral proteins Env, Tax, and p19 (Gag) were decreased in the presence of YTHDF1, while the viral protein Hbz was increased (**Figure 4C**). Accordingly, the amount of p19 (Gag) released into the supernatant was also reduced in the presence of YTHDF1 (**Figure 4D**). STM2457-treated cells depleted of m^6^A had no significant changes in transcript or protein expression in response to YTHDF1.

**Figure 4.**
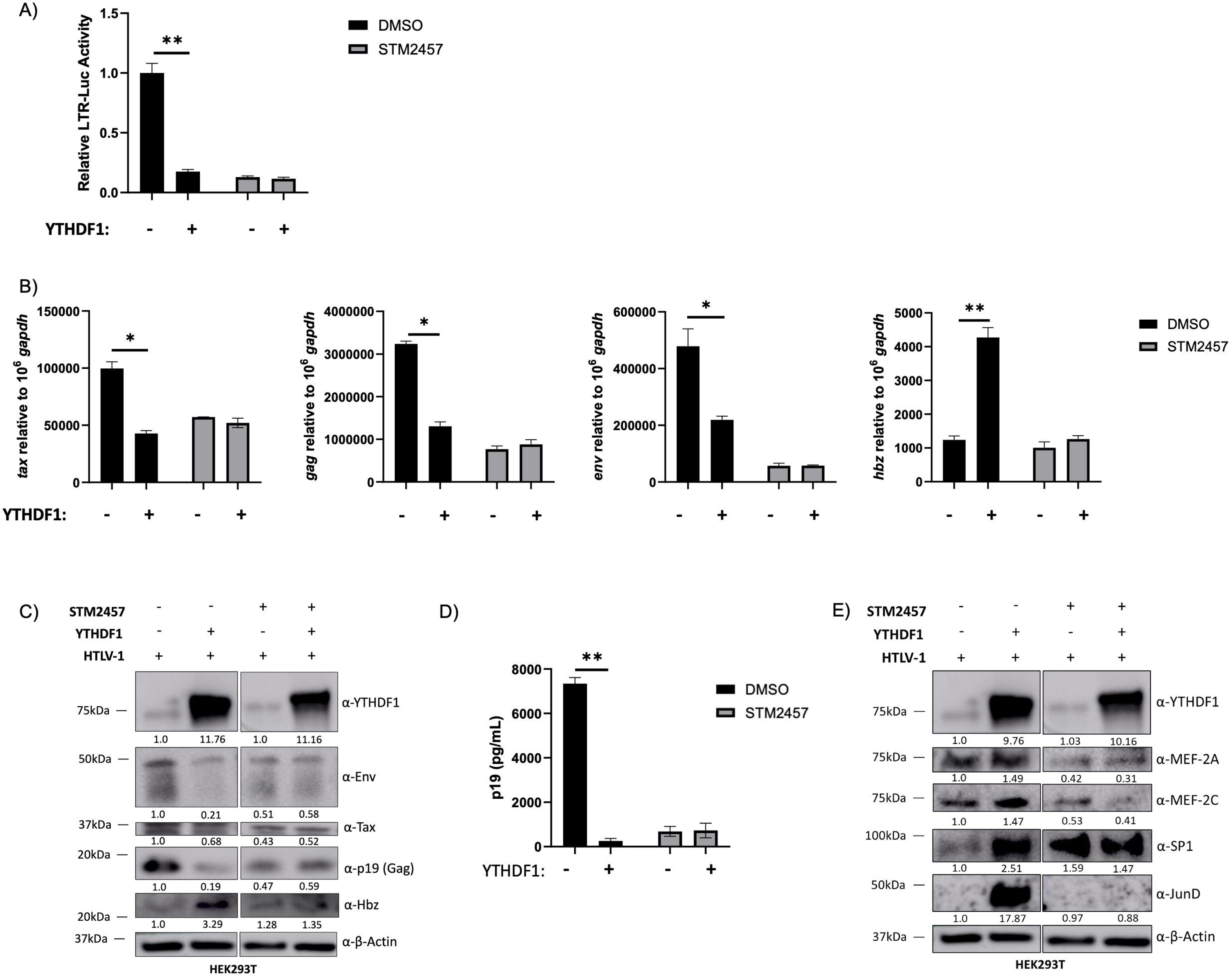
YTHDF1 decreases sense-derived and increases antisense-derived viral transcripts. HEK293T cells were treated with or without 60 μM STM2457 for 72 h and then transfected with LTR-1-luc, rtk-luc, HTLV-1 proviral DNA, and ± 100ng FLAG-YTHDF1 expression vector. Cells were collected 72 h post-transfection. A) Cells were lysed in passive lysis buffer and firefly luciferase activity was quantified relative to renilla luciferase. B) RNA was isolated and qRT-PCR was used to measure sense-derived transcripts (*tax, gag,* and *env*) and antisense-derived (*hbz*) transcripts. C) Western blot analysis was used to measure viral protein (Env, Tax, p19 [Gag], Hbz) expression. β-actin was used as a loading control. Relative band intensity was quantified using ImageJ. Protein quantification relative to no YTHDF1 (lane 1; normalized to β-actin) is depicted below each western blot. D) Virus release in the cell supernatant was measured using a p19 Gag ELISA. E) Western blot analysis was used to measure transcription factors SP1, JunD, MEF-2A, and MEF-2C. Antibodies against YTHDF1 and β-actin (loading control) were also used. Relative band intensity was quantified using ImageJ. Protein quantification relative to no YTHDF1 (lane 1; normalized to β-actin) is depicted below each western blot. Statistical significance was determined using a Student’s *t*-test; \**p* ≤ 0.05, \*\**p* ≤ 0.01.

The effect of YTHDF1 knockdown in a physiologic T-cell environment was examined in SLB-1 and ATL-ED T-cell lines. shRNA-mediated knockdown of YTHDF1 increased sense-derived viral protein and transcript levels and decreased antisense-derived Hbz protein and transcript levels (**Figure 5A, 5C, 5D, 5E**). As expected, knockdown of YTHDF1 increased viral p19 (Gag) released into the cell supernatant (**Figure 5B**). These differential effects on sense and antisense-derived viral genes were lost upon STM2457 treatment (m^6^A depletion).

**Figure 5.**
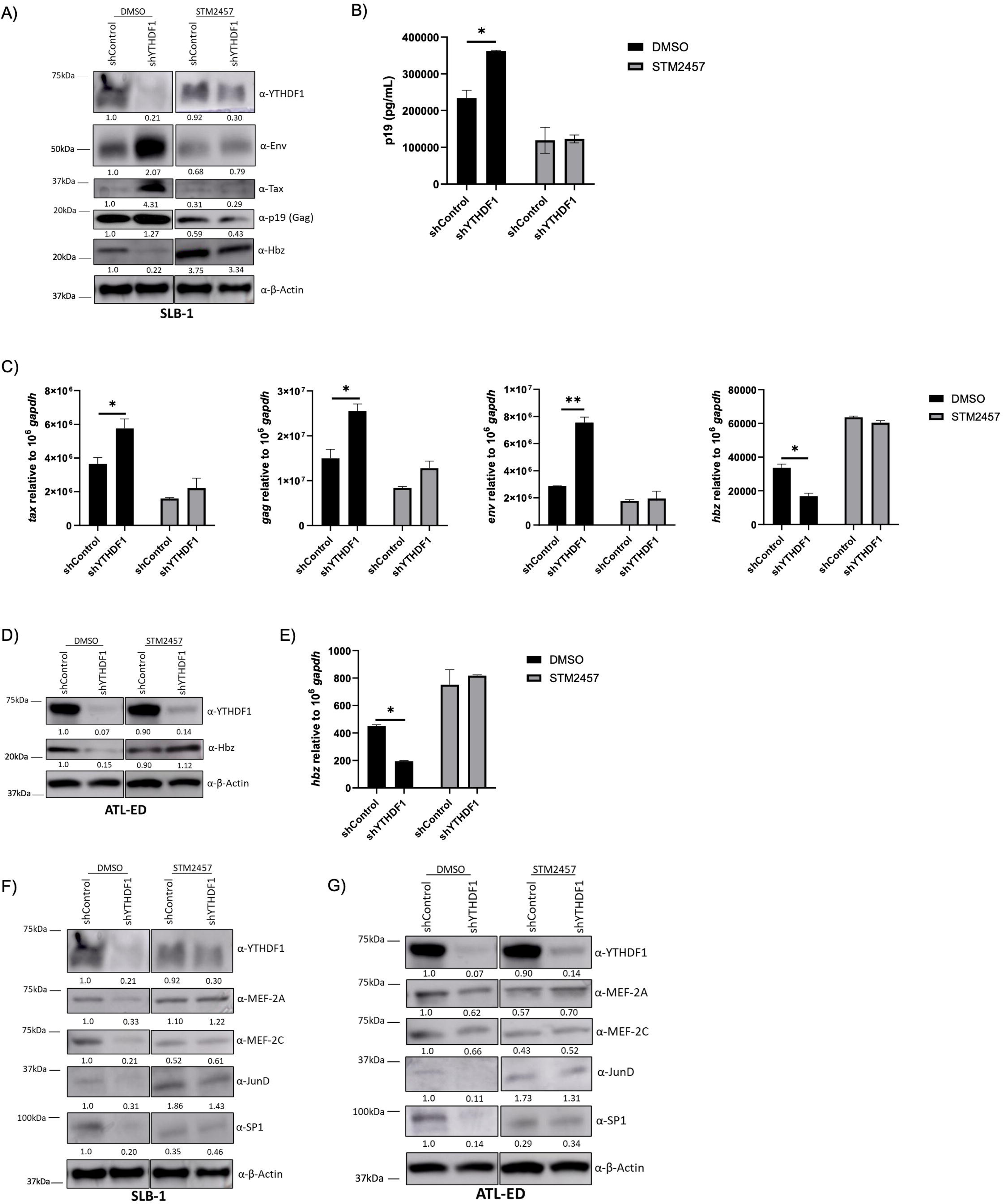
Knockdown of YTHDF1 increases sense-derived and decreases antisense-derived viral transcripts. SLB-1 and ATL-ED cells were treated with or without 60 μM STM2457 for 72 h. Cells were then transduced with shControl and shYTHDF1 lentivirus. A, D) Western blot analysis was used to measure viral protein (Env, Tax, p19 [Gag], Hbz), YTHDF1 and β-actin (loading control). Relative band intensity was quantified using ImageJ. Protein quantification relative to DMSO, shControl (lane 1; normalized to β-actin) is depicted below each western blot. B) Virus release in the cell supernatant (of SLB-1 cells) was measured using a p19 Gag ELISA. C, E) RNA was isolated and qRT-PCR was used to measure the transcript levels of *tax*, *env*, *gag*, and *hbz* relative to *gapdh.* F) SLB-1 cells and G) ATL-ED cells were subjected to immunoblot to measure YTHDF1, JunD, SP1, MEF-2A, MEF-2C, and β-actin (loading control). Relative band intensity was quantified using ImageJ. Protein quantification relative to shControl, DMSO (lane 1; normalized to β-actin) is depicted below each western blot. Statistical significance was determined using a Student’s *t*-test; \**p* ≤ 0.05, \*\**p* ≤ 0.01.

The 5’ and 3’ LTRs are activated by different subsets of transcription factors. The 5’ LTR is activated by Tax through association and recruitment of various host proteins (CREB, CBP, p300) (69). Conversely, the 3’ LTR is driven primarily by host proteins SP1, JunD, MEF-2A, and MEF-2C (70). Previous studies in acute myeloid leukemia (AML) cells found elevated m^6^A levels on *SP1* transcript which promoted its stability and translation (71). *MEF-2C* RNA has also been found to contain m^6^A modification (72). We hypothesized the effect of YTHDF1 on *hbz* expression may be due to alterations of host proteins which regulate 3’ LTR activity. Expression of YTHDF1 in HEK293T cells increased SP1, JunD, MEF-2A, and MEF-2C protein expression (**Figure 4E**). Conversely, shRNA-mediated knockdown of YTHDF1 in SLB-1 and ATL-ED cells decreased SP1, JunD, MEF-2A, and MEF-2C protein expression (**Figure 5F, 5G**). These effects were lost when cells were treated with STM2457 to deplete m^6^A. These results suggest YTHDF1 alters the expression of host factors responsible for antisense-derived *hbz* expression.

Tax serves to activate the 5’ LTR, thus promoting transcription of all sense-derived viral genes including itself. To eliminate this feed-forward loop and determine the effect of YTHDF1 on Tax in the absence of other viral genes, we measured Tax protein and transcript levels in HEK293T cells. Using an expression vector which expresses the native *tax* mRNA sequence driven by a heterologous CMV promoter, we found YTHDF1 expression decreases Tax protein (**Figure 6A**) and *tax* transcript (**Figure 6B**). This decrease in both protein and transcript is lost in the absence of m^6^A (STM2457 treatment). We next utilized a Tax expression vector with altered *tax* mRNA sequence, also driven by a heterologous CMV promoter. This altered *tax* mRNA sequence – without the alteration of Tax amino acid sequences - ablates approximately 50% of the DRACH motifs compared to the native *tax* transcript and reduced m^6^A levels on *tax* transcripts to undetectable levels (**Supplemental Figure 3A**). Using the altered *tax* expression vector, we found YTHDF1 failed to significantly decrease protein (**Figure 6C**) and transcript (**Figure 6D**) levels. Taken together, these results suggest YTHDF1 regulates *tax* transcript abundance (independent of 5’ LTR activity) through m^6^A modifications.

**Figure 6:**
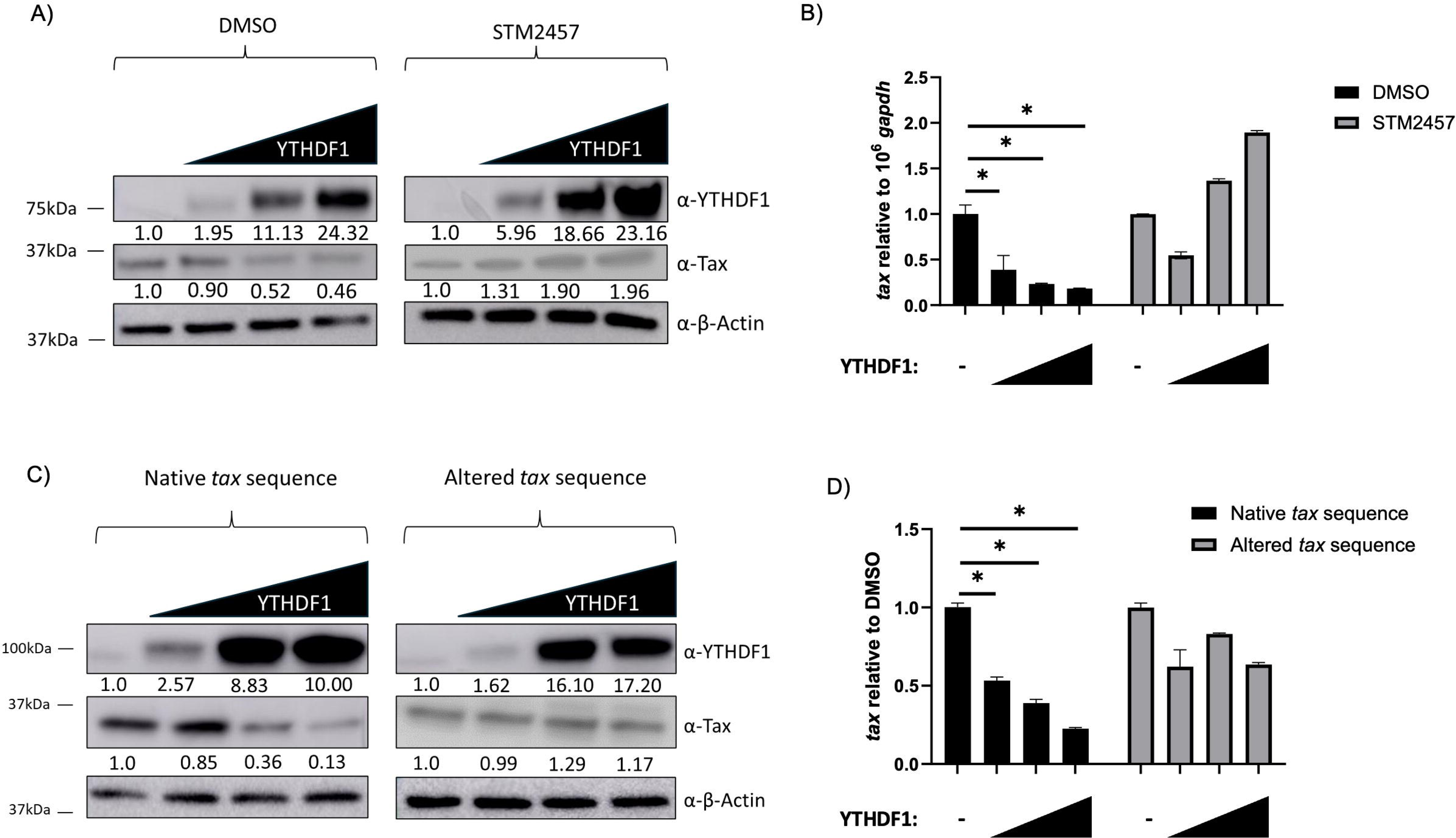
YTHDF1 decreases m^6^A-modified *tax* transcript abundance. A) HEK293T cells were treated with or without 60 μM STM2457 for 72 h and then transfected with a plasmid expressing the native *tax* RNA sequence with titrating amounts of a YTHDF1 expression vector. A) Western blot analysis was used to measure YTHDF1, Tax, and β-actin (loading control). Relative band intensity was quantified using ImageJ. Protein quantification relative to no YTHDF1 (normalized to β-actin) is depicted below each western blot. B) RNA was extracted and qRT-PCR was used to measure the transcript level of *tax* relative to *gapdh*. HEK293T cells were transfected with plasmids expressing the native *tax* RNA sequence or an altered *tax* RNA sequence with titrating amounts of a YTHDF1 expression vector. C) Western blot analysis was used to measure YTHDF1, Tax, and β-actin (loading control). Relative band intensity was quantified using ImageJ. Protein quantification relative to no YTHDF1 (normalized to β-actin) is depicted below each western blot. D) RNA was extracted and qRT-PCR was used to measure the transcript level of *tax* relative to *gapdh*. Statistical significance was determined using a Student’s *t*-test; \**p* ≤ 0.05.

### YTHDC1 increases viral transcript levels and promotes *tax* nuclear export

The functional effects of YTHDC1 on HTLV-1 biology were examined using shRNA-mediated knockdown of YTHDC1 in the leukemic SLB-1 T-cell line. Knockdown of YTHDC1 decreased viral protein Tax, Env, Gag and Hbz expression (**Figure 7A**) and p19 (Gag) release into the supernatant (**Figure 7B**). Similarly, loss of YTHDC1 decreased *tax, env, gag,* and *hbz* transcript levels (**Figure 7C**). The decrease in viral protein and transcript levels was lost when cells were treated with STM2457. Similar results were obtained in the ATLL-patient derived T-cell line, ATL- ED, which only express the *Hbz* gene (**Figure 7D, 7E**).

**Figure 7.**
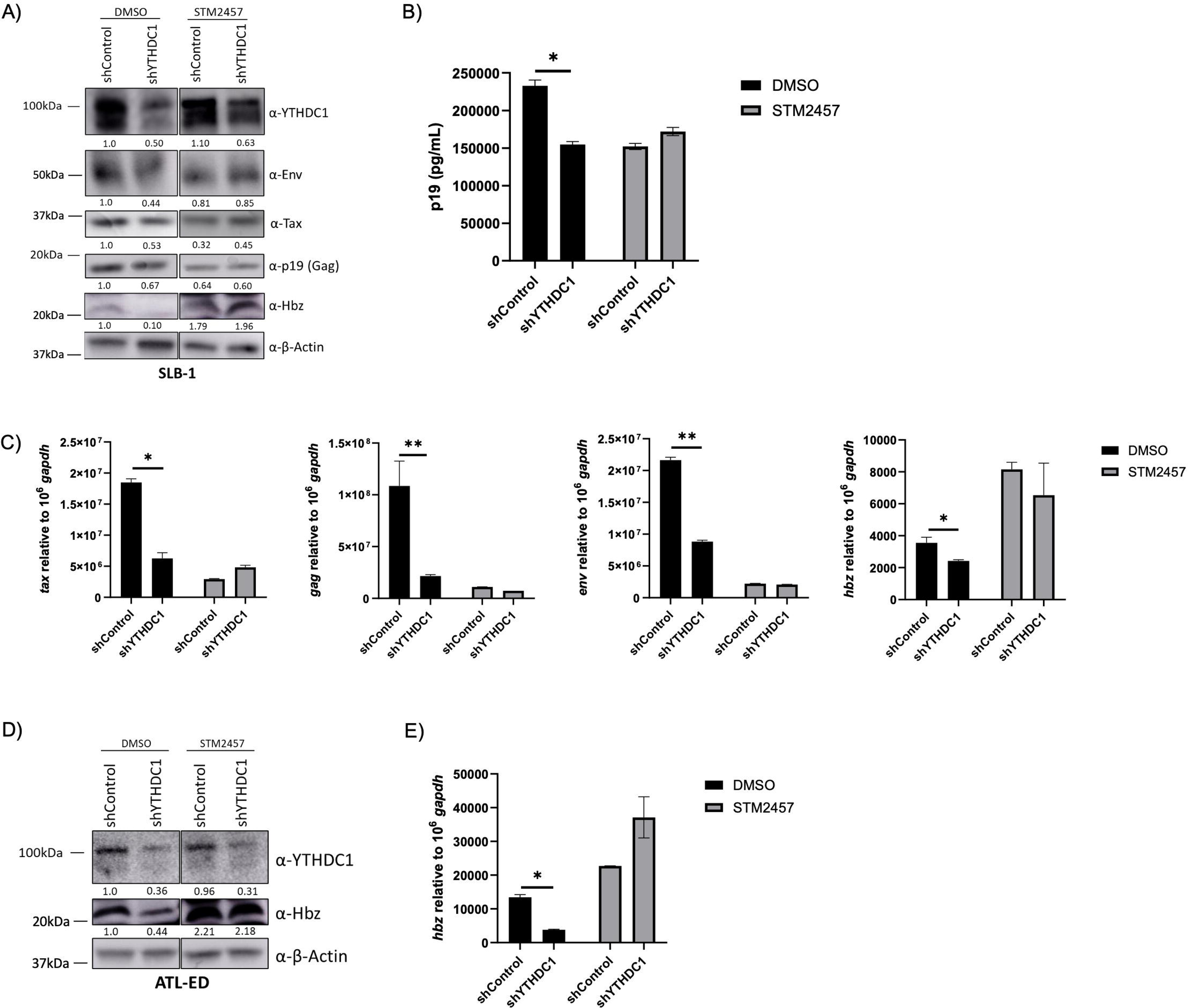
Loss of YTHDC1 decreases viral transcripts. SLB-1 and ATL-ED cells were treated with or without 60 μM STM2457 for 72 h and transduced with shControl and shYTHDF1 lentivirus. A) SLB-1 cells were subjected to western blot analysis to measure YTHDF1, Env, Tax, p19 (Gag), Hbz, and β-actin (loading control). Relative band intensity was quantified using ImageJ. Protein quantification relative to shControl, DMSO (normalized to β-actin) is depicted below each western blot. B) Virus release in the cell supernatant was measured using a p19 Gag ELISA. C) RNA was isolated and qRT-PCR was used to measure transcript levels of *tax*, *gag*, *env,* and *hbz* relative to *gapdh*. D) ATL-ED cells were subjected to western blot analysis measure YTHDF1, Hbz, and β-actin (loading control). Relative band intensity was quantified using ImageJ. Protein quantification relative to shControl, DMSO (normalized to β-actin) is depicted below each western blot. E) RNA was isolated and qRT-PCR was used to measure transcript levels of *hbz* relative to *gapdh*. Statistical significance was determined using a Student’s *t*-test; \**p* ≤ 0.05, \*\**p* ≤ 0.01.

YTHDC1 has been previously reported to regulate nuclear export of m^6^A-modified transcripts (35). We hypothesized YTHDC1 may similarly be regulating the nuclear export of m^6^A-modified viral transcripts. To investigate this possibility, vehicle (DMSO) or STM2457- treated HEK293T cells were transfected with our native *tax* expression vector with titrating levels of YTHDC1. Cells were then fractionated to measure the relative abundance of nuclear and cytoplasmic *tax* transcripts. Successful fractionation was confirmed by western blot (**Supplemental Figure 4**). Expression of YTHDC1 significantly increased the proportion of *tax* transcript in the cytoplasm (**Figure 8A**), as well as total *tax* transcript (**Figure 8B**). These alterations in *tax* transcript distribution corresponded to increased Tax protein expression (**Figure 8C**). Conversely, treatment of cells with STM2457 abolished the YTHDC1-mediated effect on *tax* localization (**Figure 8A**), total transcript abundance (**Figure 8B**) and protein expression (**Figure 8C**). Taken together, this data indicates that YTHDC1 promotes nuclear export of *tax* transcripts. Unlike the cytoplasmic reader YTHDF1, however, nuclear YTHDC1 did not exhibit any notable involvement in the regulation of differential expression between *tax* and *hbz* transcripts.

**Figure 8.**
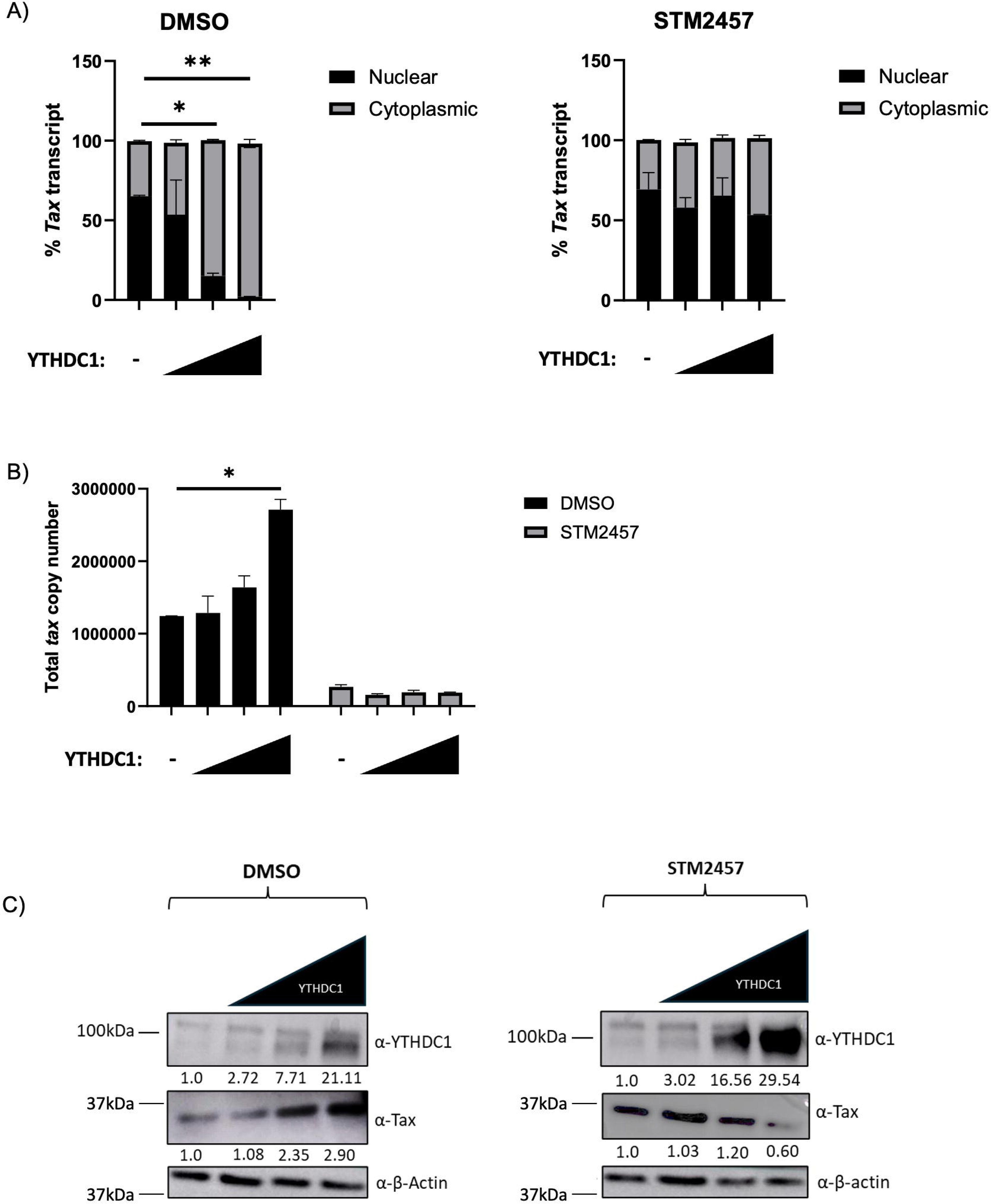
YTHDC1 promotes *tax* nuclear export. HEK293T cells were treated with or without 60 μM STM2457 for 72 h and then transfected with plasmids expressing the native *tax* RNA sequence and titrating amounts of a YTHDC1 expression vector. Cells were fractionated into nuclear and cytoplasmic extracts. RNA was isolated from each fraction using TRIzol. A) qRT- PCR was used to measure fractionated *tax* transcript levels. B) Total *tax* transcript relative to *gapdh* were measured using qRT-PCR. C) Western blot analysis of whole cell lysate was used to measure YTHDC1 and β-actin (loading control) in DMSO-treated and STM2457-treated conditions. Relative band intensity was quantified using ImageJ. Protein quantification relative to no YTHDC1 (normalized to β-actin) is depicted below each western blot. Statistical significance was determined using a Student’s *t*-test; \**p* ≤ 0.05, \*\**p* ≤ 0.01.

## Discussion

A better understanding of the mechanisms that underly HTLV-1 pathogenesis, including those that regulate *tax* and *hbz* transcripts, is critical given the severity of HTLV-1-associated diseases and ineffective treatment strategies. Epigenetic modification represents a potential regulatory mechanism of HTLV-1, which has been documented in the context of viral DNA and histones, but not viral RNA (73–76). Notably, *hbz* RNA can promote epigenetic silencing of HTLV-1 expression through interference with basal transcription machinery (77), however the RNA transcript itself has yet to be characterized as epigenetically modified.

The RNA epigenetic modification m^6^A represents an under-pursued and valuable epigenetic target in both virology and oncology. This modification has been reported in retroviruses, particularly human immunodeficiency virus type-1 (HIV-1) (78–80). HIV-1 is well documented as m^6^A-modified, with 3 major sites of m^6^A enrichment being identified: the overlap region between the env gene and the second coding exon of rev, the U3 region of the 3’ LTR, particularly in the conserved NF-κB binding sites, and the R region of the 3’ LTR coincident with the transactivation response RNA hairpin (78). Currently, conflicting data exists as to whether m^6^A modification of viral transcripts plays a promotional or inhibitory role in HIV-1 replication and infection (78, 79, 81, 82). Silencing of m^6^A writers decreased HIV-1 replication while silencing of m^6^A erasers enhanced HIV-1 replication, indicative of a positive role of m^6^A in HIV-1 replication (79). Furthermore, viral infection triggered an increase in both host and viral m^6^A-modified mRNA. These findings were supported by demonstration that YTHDF binding to m^6^A inhibits HIV-1 replication. In contrast, N’Da Konan, S *et al.* (2022) demonstrated that knockdown of METTL3/14 and YTHDF2 upregulates HIV-1 mRNA levels in infected cells (81). Tirumuru *et al.* (2016) showed that inhibition of YTHDF1-3 proteins inhibits HIV-1 infection in primary CD4+ T-cells (82). Furthermore, silencing of m^6^A writers decreased HIV-1 Gag protein expression in virus-producing cells, while silencing of m^6^A erasers increased Gag expression. Overall, these findings are indicative that while viruses, specifically retroviruses, can be m^6^A modified, there is still a limited understanding of the role of m^6^A modification in retroviral pathobiology.

Herein we present the first characterization of m^6^A in HTLV-1, describing the effects of both global m^6^A depletion and potential mechanisms of reader protein regulation of *tax* and *hbz* transcripts. We found viral RNA was m^6^A-modified and mapped m^6^A enrichment to the 3’ regulatory region of the viral RNA. In addition, the viral transcripts *hbz* and *tax* are m^6^A-modified in HTLV-1-transformed and ATLL-patient derived T-cell lines. Alteration in global m^6^A levels using the METTL3 inhibitor STM2457 demonstrated that m^6^A differentially regulates sense-derived *tax* and antisense-derived *hbz* transcripts (**Figure 9A**). The m^6^A reader proteins are responsible for carrying out the effects of m^6^A modification. We identified YTHDF1 and YTHDC1 reader proteins consistently bound *tax* and *hbz* across several different HTLV-1 cell lines. Subsequent functional studies found YTHDF1 inhibits sense-derived *tax* in part by regulating *tax* transcript abundance, whereas YTHDF1 promotes antisense-derived *hbz* transcripts by promoting the expression of host factors which drive *hbz* transcription (**Figure 9B**). YTHDC1 appears to promote viral transcript abundance in part by promoting the export of *tax* transcripts from the nucleus to the cytoplasm (**Figure 9C**).

**Figure 9.**
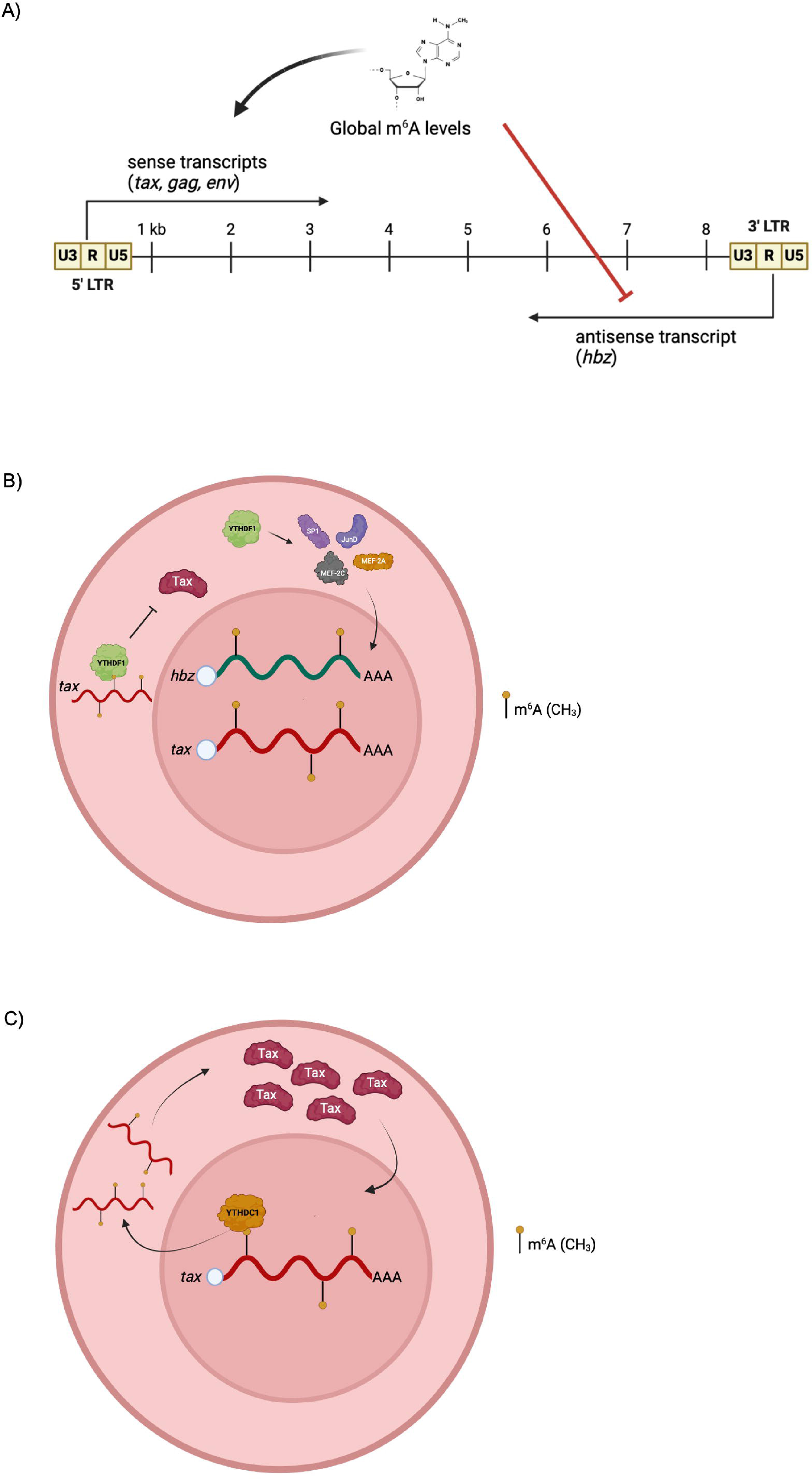
Proposed model depicting the role of global m^6^A levels, YTHDF1 and YTHDC1 in HTLV-1 infection. A) Schematic summarizing the differential regulation of *tax, gag,* and *env* (sense-derived) vs. *hbz* (antisense-derived) viral genes. The viral *tax* and *hbz* RNA transcripts are m^6^A-modified within HTLV-1-infected cells. The reader proteins YTHDF1 and YTHDC1 bind to both *tax* and *hbz*. B) YTHDF1 decreases sense-derived viral transcripts and specifically decreases *tax* transcript abundance (independent of transcription). YTHDF1 increases antisense-derived *hbz* transcript abundance at least partially through effects on transcription factors known to regulate *hbz* transcription. C) YTHDC1 increases both sense-derived and antisense-derived viral transcripts. YTHDC1 facilitates nuclear export of *tax* transcript to promote Tax protein expression.

MeRIP-Seq is used to identify m^6^A-modified RNA residues and provides a resolution of 100-200 nucleotides (23, 83). While this does not significantly change the general localization of our m^6^A peak within the genomic 3’ region, the precise nucleotides which are modified cannot be finely mapped with this technique. The imprecise single nucleotide resolution coupled with the dynamic nature of m^6^A modification likely accounts for the discrepancy in some m^6^A called peaks between replicates. Additionally, the repetitive nature of the viral LTRs somewhat distorts the ability to finely map m^6^A residues to 5’ vs. 3’ LTR repetitive sequences. Future studies which employ single nucleotide resolution mapping via Nanopore direct RNA sequencing (58, 84) in both the viral genome and viral transcriptome will be beneficial. Finely mapping site-specific m^6^A residues within HTLV-1 would allow for the evaluation of direct, site-specific m^6^A effects on viral persistence and pathogenesis. While we demonstrate other viral transcripts can be m^6^A modified, we have elected to first focus on the m^6^A post-transcriptional modification of *tax* and *hbz* given their importance in HTLV-1 pathogenesis. HTLV-1 infection and disease development are tightly controlled by the relative abundance of *tax* and *hbz*. Given that global m^6^A levels in HTLV-1-transformed cells favors increased *tax* transcripts while diminishing *hbz* transcripts, it is possible that m^6^A serves as a critical regulator of initial cellular transformation.

Although YTHDF1 and YTHDC1 reader proteins were selected for our downstream functional studies, we did observe enrichment of reader protein YTHDF3 with *tax* and *hbz* in SLB-1 cells and YTHDF2 enrichment with *hbz* in ATL-ED cells. Given that YTHDF2 is involved in mRNA decay (85), this may suggest that mRNA decay function is important for ATLL disease development. While the functional significance of YTHDF2 was not addressed in this study, future endeavors warrant focus on the role of this protein in HTLV-1 biology. The role of YTHDF3 has not been fully characterized, however is traditionally believed to have overlapping or redundant functions in regards to other reader proteins (36, 39). Recently, YTHDF3 has been noted to suppress interferon-dependent antiviral responses by promoting FOXO3 translocation (86), however this finding was not investigated in the context of HTLV-1 infection. Regardless, the documented involvement of YTHDF3 in immune regulation as it pertains to HTLV-1 supports future investigative endeavors.

This impact of YTHDF1 is notably the opposite of what we observed in cells treated with the m^6^A inhibitor STM2457. m^6^A deposition has been shown to occur co-transcriptionally (87–89), but recent insights suggest that m^6^A machinery can also impact transcription and chromatin signature (90, 91). We found YTHDF1 regulates the expression of *tax* RNA, independent of the viral 5’ LTR promoter and transcription. YTHDF1 has been shown to regulate target gene expression by promoting translation or modulating the stability of mRNAs. We found expression of YTHDF1 (either expression or reduction) affected the level of several host factors (SP1, JunD, MEF-2A, MEF-2C) responsible for activating the 3’ LTR (driving *hbz* expression). It is feasible that each of these host factors are m^6^A-modified and expression regulated by YTHDF1. Specifically, SP1 has been recently documented to have m^6^A-modified RNA in diseased renal epithelial cells (92). While the epitranscriptomic profile of SP1 has not been studied in the context of other viruses or HTLV-1, it is possible that m^6^A modification of this host factor (and potentially others) may be important in HTLV-1 disease pathogenesis.

YTHDC1 is primarily localized within the nucleus and mediates RNA fates through nuclear export, alternative splicing, RNA stabilization and RNA decay. Likewise, we found YTHDC1 increases *tax* and *hbz* transcript levels in HTLV-1 T-cell lines. Although we found YTHDC1 enhances nuclear export of *tax*, it remains unclear if the increase in Tax protein is due to increased cytoplasmic *tax* available for translation or further RNA stabilization.

While the role of m^6^A has not been evaluated in ATLL, it has been studied in the context of another blood cancer called AML. Dysregulation of the m^6^A-modifying system was initially shown to contribute to the progression of AML (93), indicating a role for m^6^A in AML pathogenesis. Importantly, inhibition of METTL3 through the use of STM2457 was found to reduce tumor burden and promote apoptosis (94). Other therapeutics have aimed at targeting the reader proteins, particularly YTHDF2, which was found to compromise AML stem cells when inactivated (95). Since our studies indicate that m^6^A and reader proteins YTHDF1 and YTHDC1 regulate the viral transcriptome (an essential phenomenon for ATLL development), subsequent chemotherapy-alternatives may be aimed at targeting these reader proteins or the m^6^A writer protein METTL3 (via STM2457).

This study presents the first description of m^6^A modification in HTLV-1 and provides a potential mechanism by which reader proteins YTHDF1 and YTHDC1 regulate *tax* and *hbz* (**Figure 9**). Future studies should further characterize the mechanisms by which all reader proteins regulate viral transcripts, given that disease development is tightly controlled by the balance between Tax and Hbz. In particular, YTHDF2 and YTHDF3 appear to bind *tax* and/or *hbz* transcripts in HTLV-1-transformed cell lines and ATLL-derived cell lines, but not in HTLV-1- immortalized cell lines. Further investigation is warranted to determine what facets of viral transformation or disease development require YTH reader proteins, and whether these proteins are inhibitory in the steps of HTLV-1-mediated T-cell immortalization. By understanding the epigenetic factors that regulate viral gene expression, protein function and interaction with the cellular environment, insight can be gained as to what allows HTLV-1 to promote disease and highlight potential m^6^A-directed therapeutic strategies.

## Acknowledgements

We thank members of the Panfil laboratory for valuable discussions and technical expertise in conducting these experiments. We thank Patrick L. Green for critical reading of this manuscript and valuable discussions.

## Author contributions

Conceptualization (ARP, EMK); Investigation (EMK, AM, KM); Formal Analysis (EMK, AM, KM, ARP); Writing – Original Draft Preparation (EMK, SK, ARP); Writing – Review and Editing (all)

