## Supplemental Figures for "YTHDF1 and YTHDC1 m^6^A reader proteins regulate HTLV-1 *tax* and *hbz* activity"

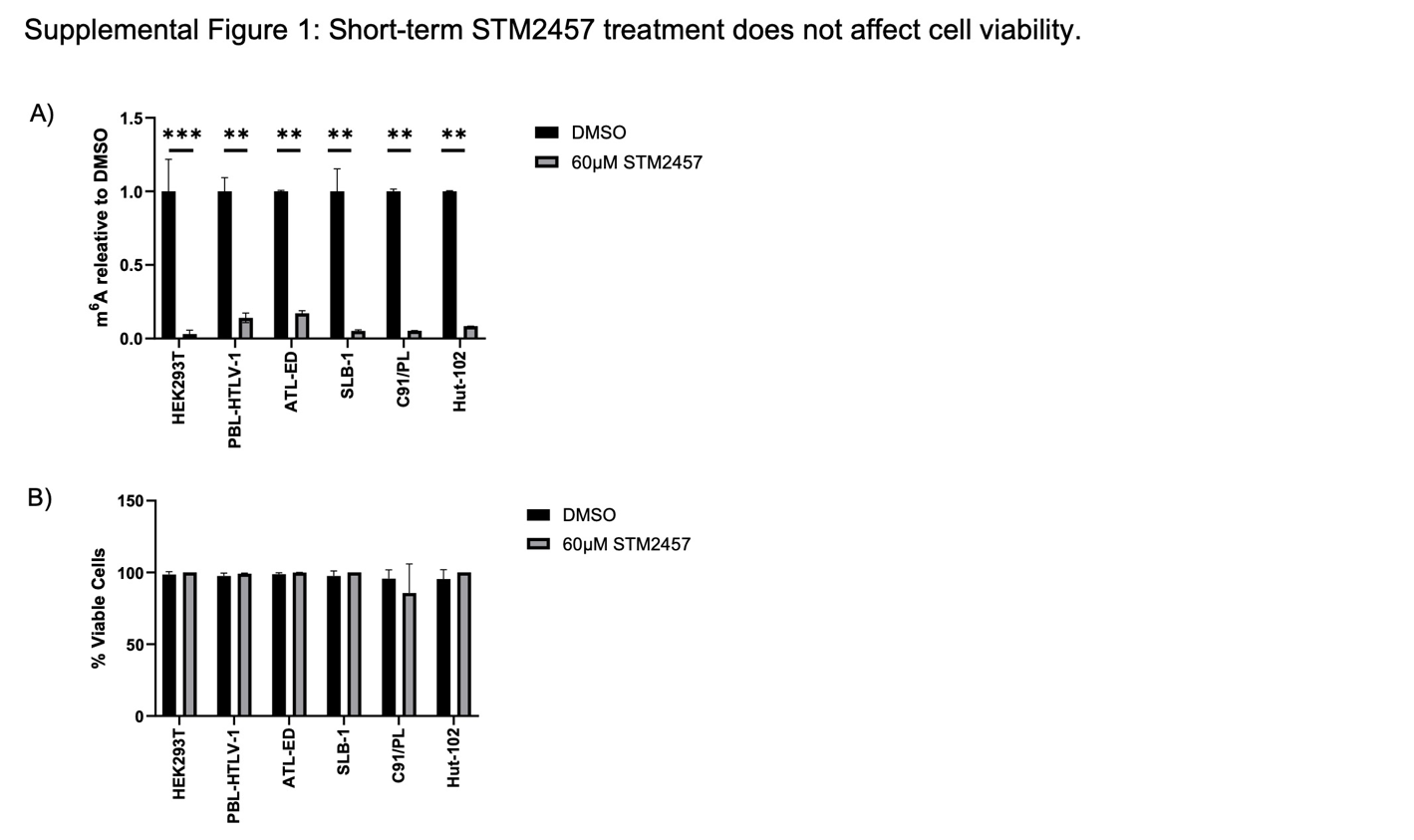
**Supplemental Figure 1. Short-term STM2457 treatment does not affect cell viability.** HEK293T, PBL-HTLV-1, ATL-ED, SLB-1, C91/PL and Hut-102 cells were treated with or without 60 μM STM2457 for 72. A) The level of m^6^A-modified mRNA in the cell was quantified in DMSO and STM2457-treated conditions using an ELISA. B) Cells were stained with 1X trypan blue and enumerated by light microscopy. Statistical significance was determined using a Student’s *t*-test; ***p* ≤ 0.01, ****p* ≤ 0.001.

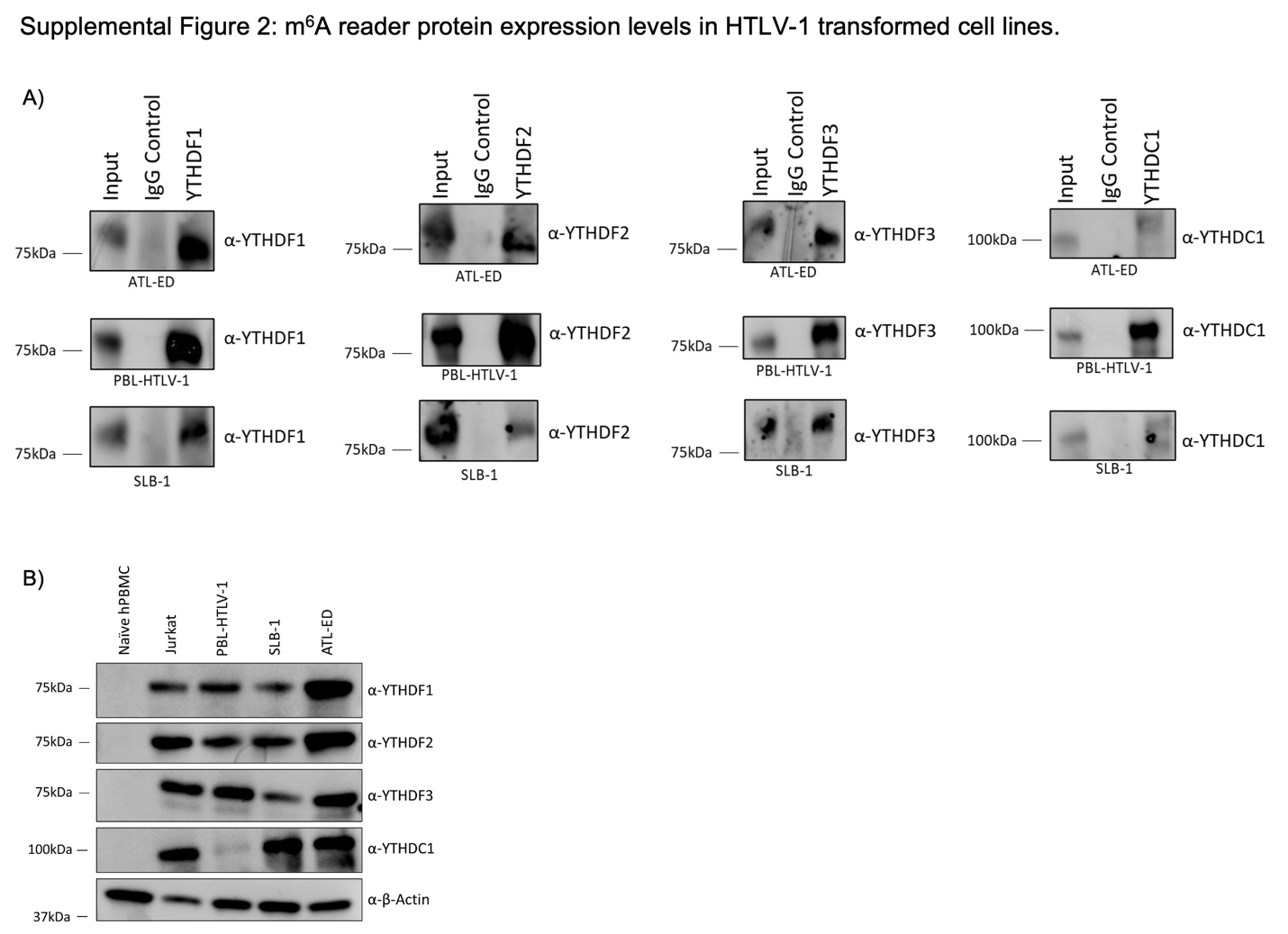

**Supplemental Figure 2. m^6^A reader protein expression levels in HTLV-1-transformed cell lines.** ATL-ED, PBL-HTLV-1 and SLB-1 cells were subjected to RNA cross-linking and immunoprecipitation using antibodies against YTHDF1-3, YTHDC1 and an IgG control. A) Western blot analysis was performed using protein lysate to ensure successful immunoprecipitation of YTHDF1, YTHDF2, YTHDF3 and YTHDC1. Whole cell lysates were applied to western blot to measure YTHDF1, YTHDF2, YTHDF3, YTHDC1 and β-actin (loading control).

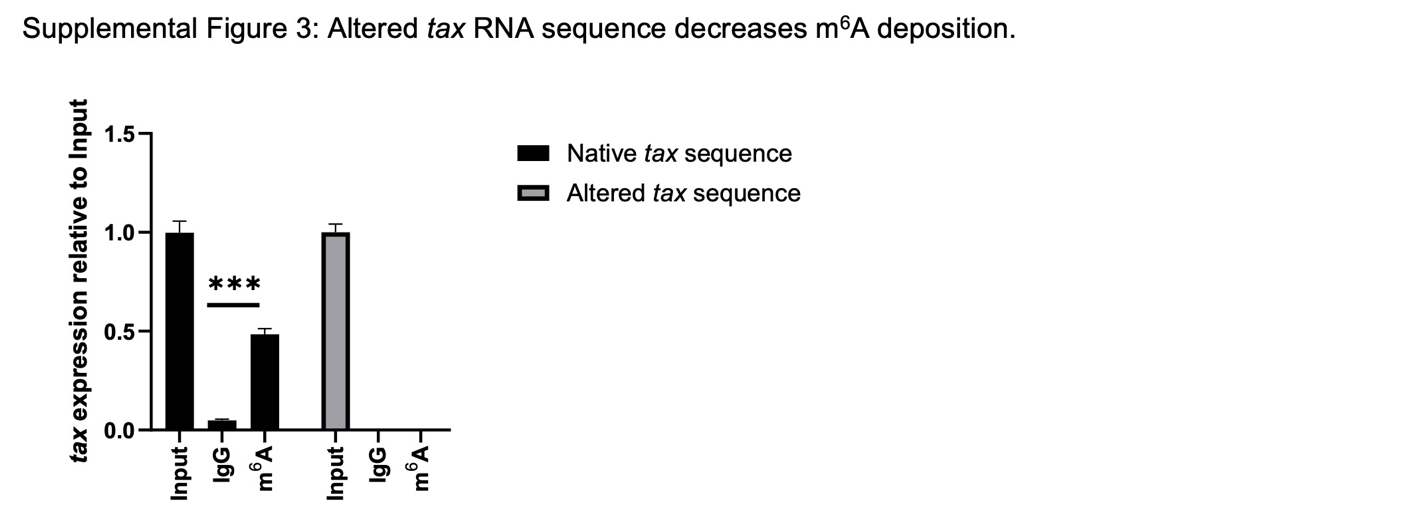

**Supplemental Figure 3. Altered *tax* RNA sequence decreases m^6^A deposition.** HEK293T cells were transfected with plasmids expressing the native *tax* sequence or an altered *tax* sequence. Cells were immunoprecipitated using antibodies against m^6^A and IgG control. RNA was isolated by TRIzol extraction and qRT-PCR was used to measure *tax* abundance relative to *gapdh*. Statistical significance was determined using a Student’s *t*-test; ****p* ≤ 0.001.

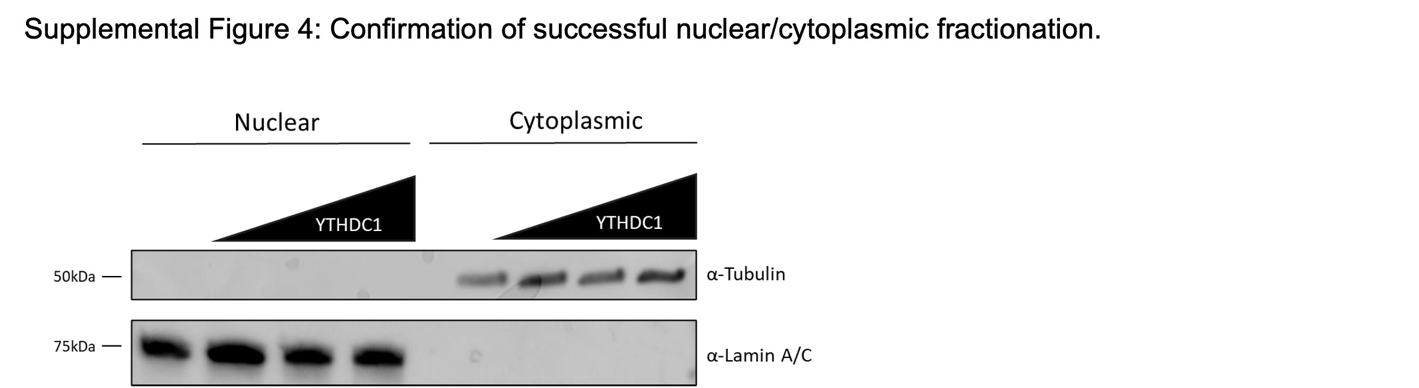

**Supplemental Figure 4. Confirmation of successful nuclear/cytoplasmic fractionation.** HEK293T cells were transfected with plasmids expressing the native *tax* sequence and increasing amounts of a YTHDC1 expression vector. Cells were fractionated into nuclear and cytoplasmic extracts. Protein lysate obtained from fractionated samples was applied to a western blot to measure abundance of β-tubulin (cytoplasmic) and Lamin A/C (nuclear) protein.
